# Myogenic Anti-Nucleolin Aptamer iSN04 Inhibits Proliferation and Promotes Differentiation of Vascular Smooth Muscle Cells

**DOI:** 10.1101/2024.04.11.588987

**Authors:** Mana Miyoshi, Takeshi Shimosato, Tomohide Takaya

## Abstract

De-differentiation and subsequent increased proliferation and inflammation of vascular smooth muscle cells (VSMCs) is one of the mechanisms of atherogenesis. Maintaining VSMCs in a contractile differentiated state is therefore a promising therapeutic strategy for atherosclerosis. We have reported the 18-base myogenetic oligodeoxynucleotide, iSN04, which serves as an anti-nucleolin aptamer and promotes skeletal and myocardial differentiation. The present study investigated the effect of iSN04 on VSMCs because nucleolin has been reported to contribute to VSMC de-differentiation under pathophysiological conditions. Nucleolin was localized in the nucleoplasm and nucleoli of both rat and human VSMCs. iSN04 without carrier was spontaneously incorporated into VSMCs, indicating that iSN04 would serve as an anti-nucleolin aptamer. iSN04 treatment decreased the ratio of EdU^+^ proliferating VSMCs and increased the expression of α-smooth muscle actin, a contractile marker of VSMCs. iSN04 also suppressed angiogenesis of mouse aortic rings ex vivo, which is a model of pathological angiogenesis involved in plaque formation, growth, and rupture. These results demonstrate that antagonizing nucleolin with iSN04 preserves VSMC differentiation, providing a nucleic acid drug candidate for the treatment of vascular disease.

## 1. Introduction

Atherosclerosis is a major risk factor for cardiovascular disease, the leading cause of death worldwide [1]. Vascular smooth muscle cells (VSMCs) in the arterial media play a key role in the development and rupture of atherosclerotic plaques [2]. Unlike skeletal and cardiac muscle cells, smooth muscle cells (SMCs), including VSMCs, can plastically switch their phenotypes of proliferation and differentiation. In the normal state, differentiated VSMCs express SMC markers such as α-smooth muscle actin (α-SMA), transgelin (SM22α), and caldesmon to maintain their contractile properties required for blood vessels [3]. However, during atherogenesis, VSMCs de-differentiate, reduce levels of SMC markers, and increase their capacity for migration, proliferation, and inflammation. This phenotypic transition causes VSMCs to migrate from the media to the intima, and the modulated VSMCs proliferate and further transform into macrophage-like cells within atherosclerotic lesions. Eventually, inflammation, apoptosis, and extracellular matrix breakdown of the de-differentiated VSMCs lead to plaque rupture, resulting in myocardial infarction or stroke [2]. Note that de-differentiation and proliferation of VSMCs occur not only in atherosclerosis, but also in restenosis, calcification, and pathological angiogenesis [4]. Therefore, maintaining VSMCs in a contractile differentiated state is an important strategy for prevention and therapy of vascular diseases.

We have reported a series of myogenetic oligodeoxynucleotides (myoDNs), which are 12-18-base single-stranded oligodeoxynucleotides that promote myogenesis [5,6]. One of the myoDNs, iSN04 (5’-AGA TTA GGG TGA GGG TGA-3’), serves as an anti-nucleolin aptamer and induces myogenic differentiation of myoblasts [5-9] and rhabdomyosarcoma cells [10]. Interestingly, iSN04 facilitates myocardial differentiation of pluripotent stem cells [11], suggesting that nucleolin antagonism by iSN04 affects not only skeletal muscle but also multiple muscle lineages. Indeed, iSN04 exerted anti-inflammatory effects on both skeletal and smooth muscle cells [12]. These studies demonstrate that iSN04 is a potential nucleic acid drug for skeletal, cardiac, and smooth muscle dysfunction, but its action on VSMC differentiation has not been investigated.

The target of iSN04, nucleolin, is a multifunctional phosphoprotein that is ubiquitously expressed and changes its subcellular localization depending on biological processes such as proliferation, differentiation, and inflammation [13]. Nucleolin has been reported to localize to the nucleoli of rat aortic SMCs [14]. In VSMCs, angiotensin II induces nucleolin expression and its translocation from the nucleus to the cytoplasm. Cytoplasmic nucleolin binds and stabilizes epidermal growth factor (EGF) and platelet-derived growth factor (PDGF) mRNAs, leading to a phenotypic transformation that decreases SMC markers and increases proliferation [15,16]. In *ApoE*^-/-^ mice fed with high-fat diet, nucleolin expression is upregulated in VSMCs within advanced aortic plaques. An atherogenic cholesterol, oxidized low-density lipoprotein (oxLDL), increases nucleolin, which is predicted to interact with aurora B to promote the cell cycle [17]. Nucleolin is also involved in oxLDL-induced foam cell formation [18]. In pulmonary arterial SMCs, hypoxia induces nucleolin translocation from the nucleus to the plasma membrane. Neurite growth-promoting factor 2 (midkine) interacts with cell surface nucleolin and activates EGF receptor signaling to promote migration and proliferation [19]. These studies reproducibly present that nucleolin contributes to the de-differentiation and proliferation of VSMCs under pathophysiological conditions, and strongly suggest that antagonizing nucleolin would be beneficial for VSMC treatment.

The present study investigated the effect of the myogenetic anti-nucleolin aptamer, iSN04, on the proliferation and differentiation of VSMCs in vitro and on the angiogenesis of aortic rings ex vivo. As expected iSN04 maintained VSMC differentiation, providing a feasible scheme for atherosclerosis therapy.

## 2. Materials and Methods

### 2.1. Chemicals

iSN04 in which all phosphodiester bonds were phosphorothioated to enhance nuclease resistance in cell culture was synthesized and HPLC-purified (GeneDesign, Osaka, Japan). 6-FAM-iSN04 is the iSN04 conjugated to 6-carboxyfluorecein at the 5’ end (gene Design) [5,20].

### 2.2. Cell Culture

All cells were cultured at 37°C under 5% CO_2_ throughout the experiments. Embryonic rat thoracic aortic SMC line A10 (CRL-1476; ATCC, Manassas, VA, USA) was maintained in growth medium (GM) consisting of DMEM (Nacalai, Osaka, Japan), 10% fetal bovine serum (FBS) (GE Healthcare, Chicago, IL, USA), and a mixture of 100 units/ml penicillin and 100 μg/ml of streptomycin (P/S) (Nacalai). A10 cells were induced to differentiate in differentiation medium (DM) consisting of DMEM, 2% horse serum (HyClone; Cytiva, Marlborough, MA, USA), and P/S.

The commercially available human aortic SMC (hAoSMC) stocks from healthy 55-year-old male and 51-year-old female (C-12533, lot 437Z012.2 and 437Z016.2, respectively; PromoCell, Heidelberg, Germany) were maintained in Smooth Muscle Cell Growth Medium 2 Kit (C-22162; PromoCell) as GM for hAoSMC (hGM) and induced to differentiate in DM for hAoSMC (hDM) consisting of DMEM, 1% FBS, and P/S.

### 2.3. Immunocytochemistry

1.0×10^5^ A10 cells and 7.5×10^4^ hAoSMCs were seeded on 30-mm dishes. For α-SMA staining, A10 cells were treated the next day with 3 or 10 μM iSN04 in DM for 4 days. The cells were fixed with 2% paraformaldehyde, permeabilized with 0.2% Triton X-100, and immunostained with 1.0 μg/ml rabbit polyclonal anti-nucleolin antibody (ab22758; Abcam, Cambridge, UK) or 1.0 μg/ml mouse monoclonal anti-α-SMA antibody (1A4, ab7817; Abcam) overnight at 4°C and then with 0.1 μg/ml Alexa Fluor 594-or 488-conjugated donkey polyclonal anti-rabbit or anti-mouse IgG antibody (Jackson ImmunoResearch, West Grove, PA, USA) for 1 h at room temperature. Cell nuclei were stained with DAPI (Nacalai). Fluorescence images were captured using EVOS FL Auto microscope (AMAFD1000; Thermo Fisher Scientific, Waltham, MA, USA). SMA signal intensity was quantified using ImageJ software (National Institutes of Health, Bethesda, MD, USA).

### 2.4. Incorporation Assay

1.0×10^4^ A10 cells/well were seeded on glass chamber slides (SCS-N08; Matsunami Glass, Osaka, Japan) and treated the next day with 5 μg/ml 6-FAM-iSN04 in GM for 0.5-4 h. The cells were then washed with PBS, fixed with 2% paraformaldehyde, and stained with DAPI. Fluorescence images were captured using EVOS FL Auto microscope [5,20].

### 2.5. EdU Staining

1.0×10^5^ A10 cells or 9.0×10^4^ hAoSMCs were seeded on 30-mm dishes and treated the next day with 30 μM iSN04 in GM for 24 h (A10) or 48 h (hAoSMC). EdU (5-ethynyl-2’-deoxyuridine) was administered at a final concentration of 10 μM, and the cells were cultured for 3 h (A10) or 12 h (hAoSMC). EdU staining was performed using the Click-iT EdU Imaging Kit (Thermo Fisher Scientific), according to the manufacturer’s instruction. Cell nuclei were stained with DAPI. Fluorescent images were captured using EVOS FL Auto microscope. The ratio of EdU^+^ cells was defined as the number of EdU^+^ nuclei divided by the total number of nuclei using ImageJ software [10].

### 2.6. Quantitative Real-Time RT-PCR (qPCR)

1.0×10^5^ hAoSMCs were seeded on 60-mm dishes and treated the next day with 30 μM iSN04 in hGM for 48 h. Total RNA was isolated using NucleoSpin RNA Plus (Macherey-Nagel, Düren, Germany) and reverse transcribed using ReverTra Ace qPCR RT Master Mix (TOYOBO, Osaka, Japan). qPCR was performed using GoTaq qPCR Master Mix (Promega, Madison, WI, USA) with the StepOne Real-Time PCR System (Thermo Fisher Scientific). The amount of each transcript was normalized to that of 3-monooxygenase/tryptophan 5-monooxygenase activation protein zeta gene (*YWHAZ*). The results are presented as fold-changes. The primer sequences of *ACTA2* [21], *CALD1* [22], *MKI67* [23], *TAGLN* [21], and *YWHAZ* [5] have been described previously.

### 2.7. Aortic Ring Assay

Three-dimensional aortic ring angiogenesis was analyzed as previously reported [24]. Aortas from 9-week-old male C57BL/6J mice (Japan SLC, Shizuoka, Japan) were dissected, perfused with Opti-MEM (Thermo Fisher Scientific), cut into 1-mm rings, and cultured in Opti-MEM for 24 h. The serum-starved aortic rings were placed in 50 μl collagen gel solution, consisting of 80% Cellmatrix Type I-A (Nitta Gelatin, Osaka, Japan), 10% 10×DMEM (Sigma-Aldrich, St. Louis, MO, USA), 0.22% NaHCO_3_, 20 mM HEPES, and 5 mM NaOH, which was pre-seeded in 96-well plates on ice. The plates were then incubated at 37°C for 30 min. Gel-embedded aortic rings were cultured in 100 μl/well GM for 6 days and captured using EVOS FL Auto microscope. Angiogenesis was quantified using ImageJ software. The ends of neovessels were joined to outline the total area. Neovascular sprouting area was calculated as the difference between the number of pixels of the total area and the aortic area defined by the circumference of the ring [25].

### 2.8. Statistical Analysis

The results are presented as the mean ± standard error. Statistical comparisons between two groups were performed using the unpaired two-tailed Student’s *t*-test and among multiple groups using Scheffe’s *F* test following one-way analysis of variance. Statistical significance was set at *p* < 0.05.

## 3. Results

### 3.1. Nucleolin Localization and iSN04 Incorporation in A10 Cells

The effect of iSN04 on VSMCs was investigated using the embryonic rat thoracic aortic SMC line A10, which resembles de-differentiated SMCs and neointimal cells [26]. Immunostaining revealed that nucleolin is expressed and localized in the nuclei, particularly in the nucleoli, of proliferating A10 cells in GM (Figure 1A, day 0), as previously reported in rat aortic SMCs [14]. Nucleolin localization was not altered during differentiation induced by DM for 4 days regardless of iSN04 treatment (Figure 1A, day 4). iSN04 must be taken up into the cytoplasm and then into the nucleoplasm to reach nuclear nucleolin. Intracellular uptake of iSN04 without a carrier has been reported in myoblasts [5]. As shown in Figure 1B, 6-FAM-iSN04 was autonomously internalized into A10 cells within 30 min and diffused into the cytoplasm and nucleoplasm by 4 h, which is generally observed for oligodeoxynucleotide transport by gymnosis [5,20,27]. These results demonstrate that iSN04 as an anti-nucleolin aptamer is able to act intracellularly in A10 cells.

**Figure 1.**
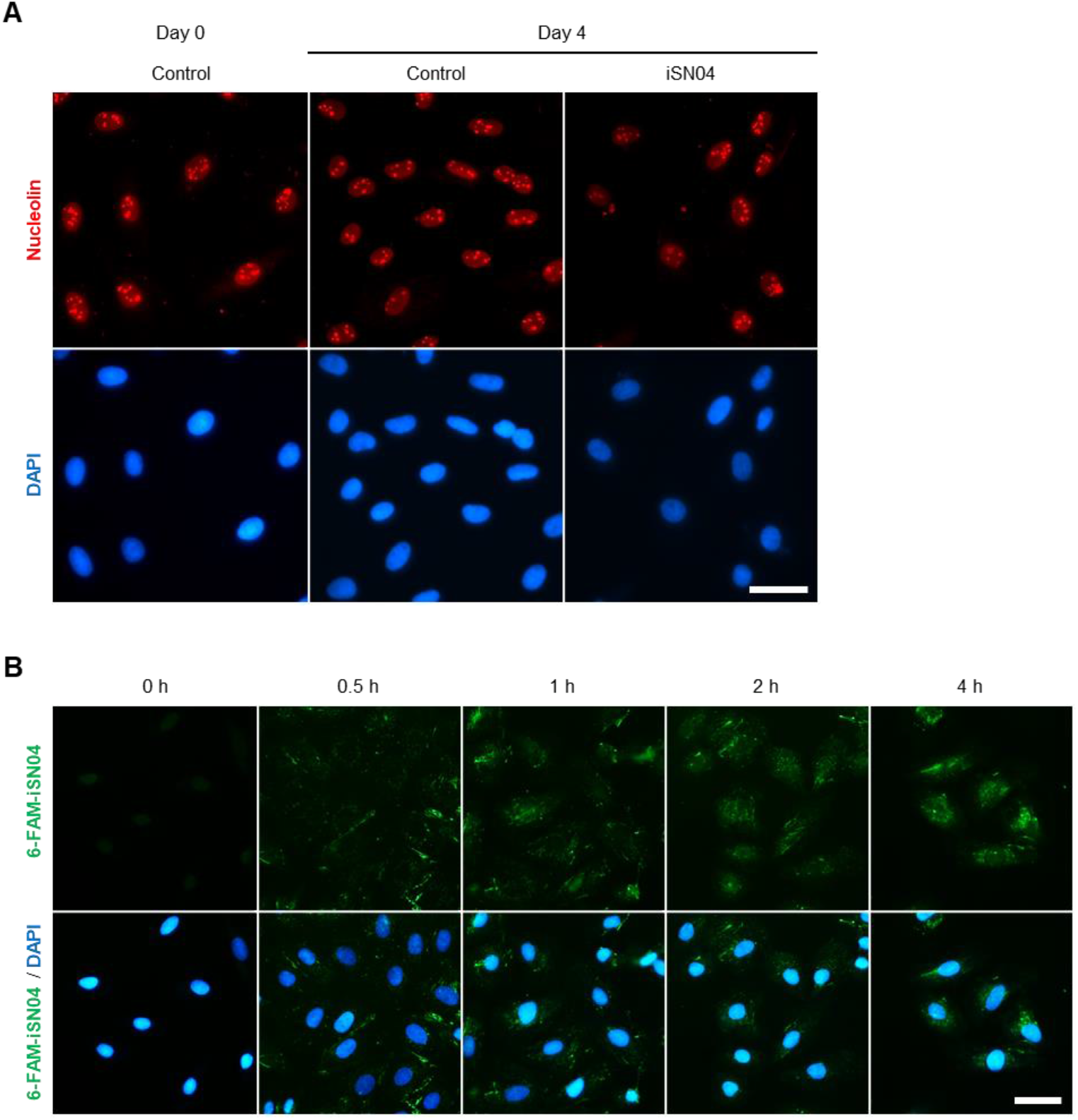
Nucleolin localization and iSN04 incorporation in A10 cells. (**A**) Representative fluorescence images of nucleolin staining of A10 cells in GM (day 0) and DM with or without 10 μM iSN04 (day 4). Scale bar, 50 μm. (**B**) Representative fluorescence images of A10 cells treated with 5 μg/ml 6-FAM-iSN04 in GM. Scale bar, 50 μm.

### 3.2. iSN04 Suppresses Proliferation and Promotes Differentiation of A10 Cells

To verify the effect of iSN04 on proliferation, A10 cells were subjected to EdU staining. iSN04 treatment significantly decreased the ratio of EdU^+^ cells replicating genomic DNA (Figure 2A), indicating that iSN04 delayed the cell cycle. Next, the effect of iSN04 on A10 cell differentiation was quantified by α-SMA staining. As shown in Figure 2B, iSN04 significantly decreased α-SMA signal intensities per cell in a dose-dependent manner during the induction of differentiation. These data demonstrate that antagonizing nucleolin with iSN04 suppresses proliferation and promotes differentiation of A10 cells.

**Figure 2.**
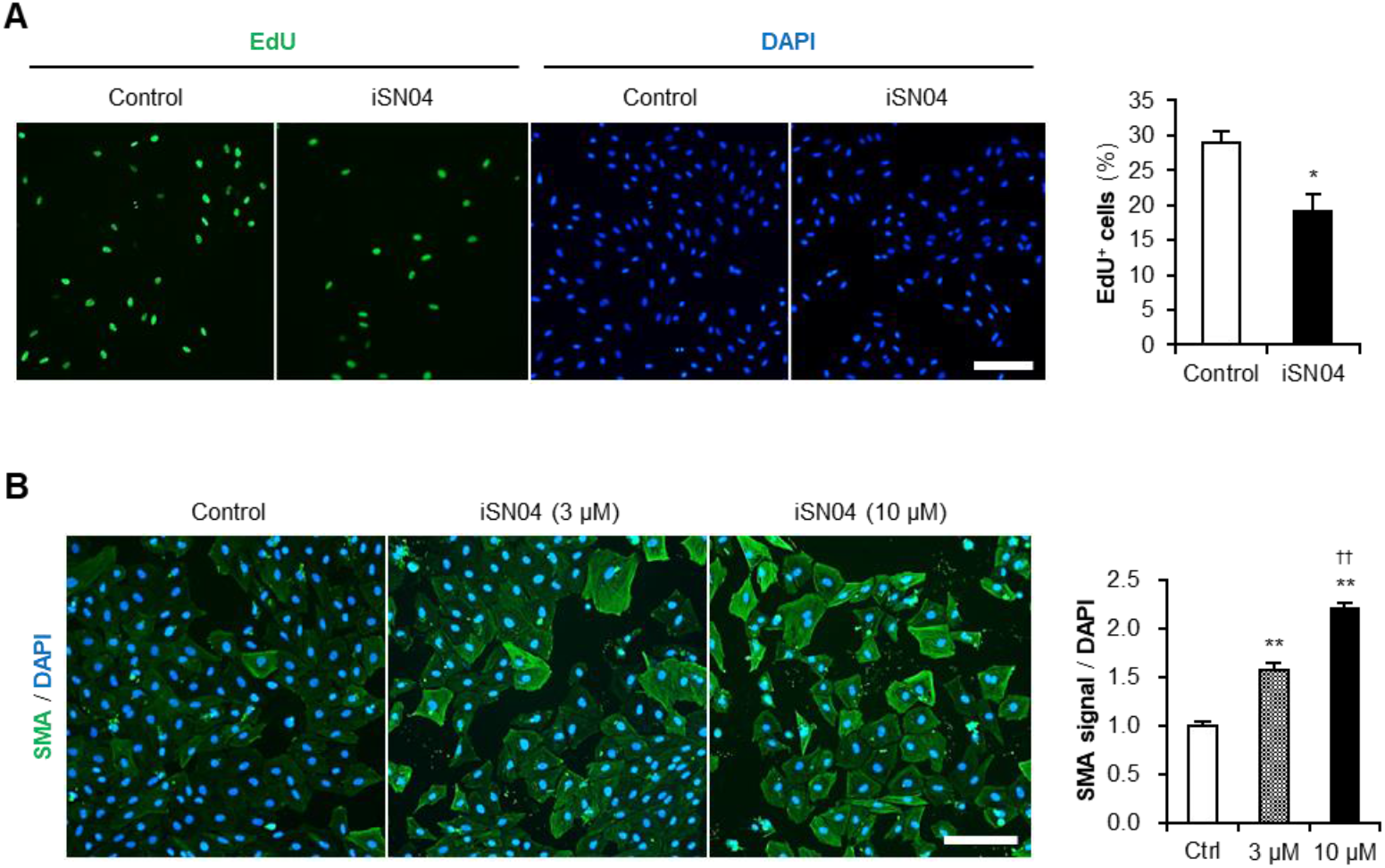
The Effect of iSN04 on proliferation and differentiation of A10 cells. (**A**) Representative fluorescence images of EdU staining of A10 cells pre-treated with 30 μM iSN04 in GM for 24 h and then with 10 μM EdU in GM for 3 h. Scale bar, 200 μm. The ratio of EdU^+^ cells were quantified. * *p* < 0.05 vs control (Student’s *t*-test). *n* = 4. (**B**) Representative fluorescence images of α-SMA staining of A10 cells treated with 3 or 10 μM iSN04 in DM for 4 days. Scale bar, 200 μm.α-SMA signal intensities per number of DAPI^+^ nuclei were quantified. ** *p* < 0.01 vs control, ^††^ *p* < 0.01 vs 3 μM iSN04 (Scheffe’s *F* test). *n* = 7.

### 3.3. iSN04 Suppresses Proliferation and Promotes Differentiation of hAoSMCs

To evaluate the effect of iSN04 on human VSMCs, primary-cultured hAoSMC stocks were used. As shown in Figure 3A, immunostaining revealed that nucleolin is expressed and localized in the nucleoplasm and nucleoli of hAoSMCs as observed in A10 cells in both proliferating and differentiating conditions. Since iSN04 promotes myogenesis of both murine and human myoblasts [5], it assumed that iSN04 would not only affect rat A10 cells but also hAoSMCs. EdU staining showed that iSN04 significantly suppressed the growth of hAoSMCs (Figure 3B). Correspondingly, qPCR results indicated that iSN04 treatment significantly decreased the mRNA level of Ki-67 (*MKI67*), a cell proliferation marker (Figure 3C). While the expression of contractile SMC markers, α-SMA (*ACTA2*), SM22α (*TAGLN*), and caldesmon (*CALD1*), were significantly upregulated by iSN04 (Figure 3C). These data clearly demonstrate that iSN04 suppresses proliferation and induces differentiation of human VSMCs, which may be applicable for clinical use.

**Figure 3.**
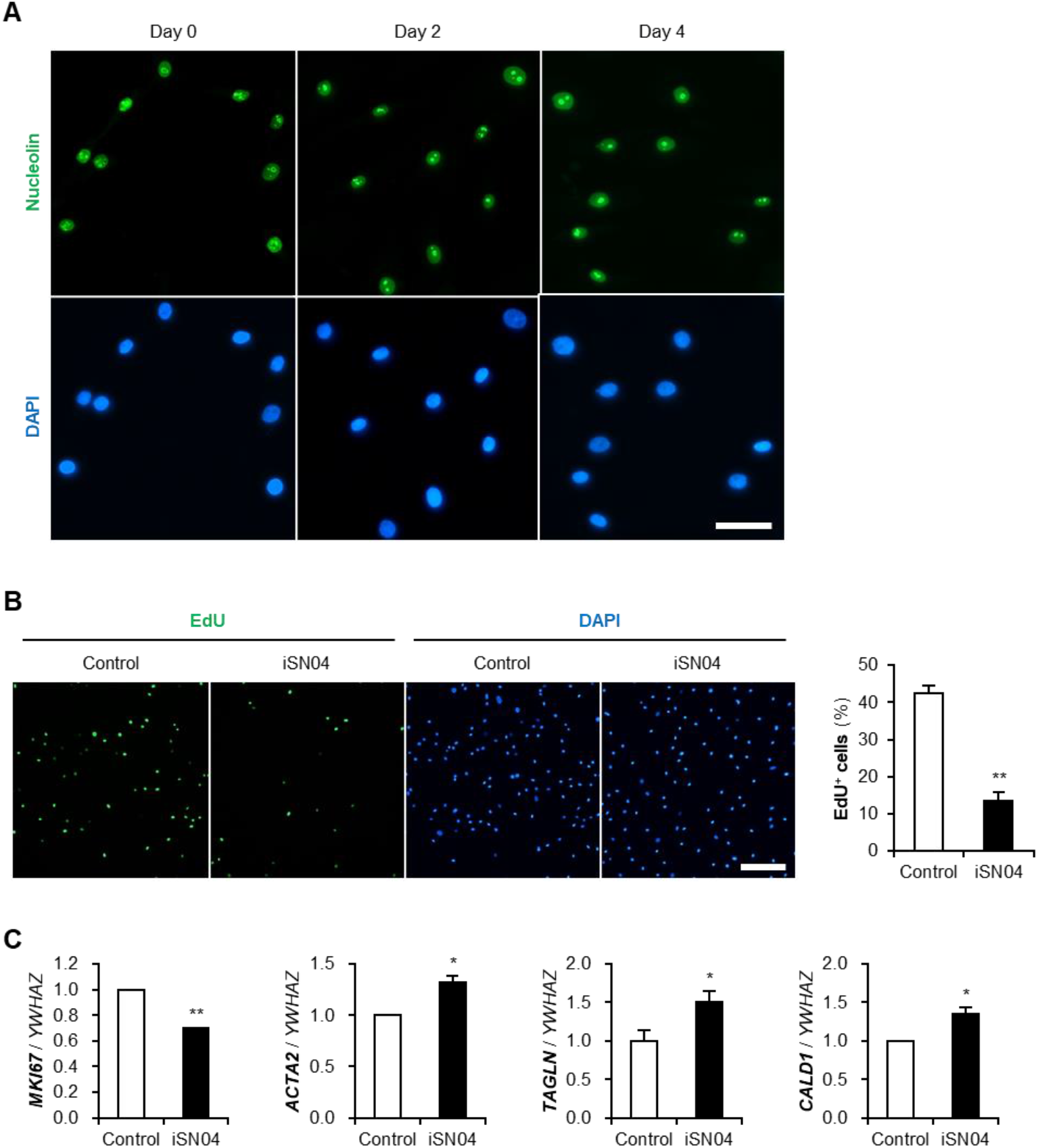
The effect of iSN04 on proliferation and differentiation of hAoSMCs. (**A**) Representative fluorescence images of nucleolin staining of hAoSMCs in hGM (day 0) and hDM (day 4). Scale bar, 50 μm. (**B**) Representative fluorescence images of EdU staining of hAoSMCs pre-treated with 30 μM iSN04 in hGM for 48 h and then with 10 μM EdU in hGM for 12 h. Scale bar, 200 μm. The ratio of EdU^+^ cells were quantified. ** *p* < 0.01 vs control (Student’s *t*-test). *n* = 4. (**C**) qPCR results of cell-cycle and SMC marker genes in hAoSMCs treated with 30 μM iSN04 in hGM for 48 h. * *p* <0.05, ** *p* < 0.01 vs control (Student’s *t*-test). *n* = 3.

### 3.4. iSN04 Suppresses Aortic Ring Angiogenesis

Phenotype switching of VSMCs is a mechanism of pathological angiogenesis that contributes to atherosclerotic plaque formation, growth, and rupture [28,29]. Since inhibition of angiogenesis renders atherosclerotic lesions small and stable, it may be a potential therapeutic strategy [29]. The aortic ring assay is a three-dimensional model of pathophysiological microvessel formation to test the anti-angiogenic effect of molecules ex vivo [24]. As shown in Figure 4, mouse aortic rings were embedded in collagen gels and cultured with or without iSN04 for 6 days. The neovascular sprouting area was significantly reduced by iSN04 treatment, indicating that iSN04 suppresses neoangiogenesis in the aorta.

**Figure 4.**
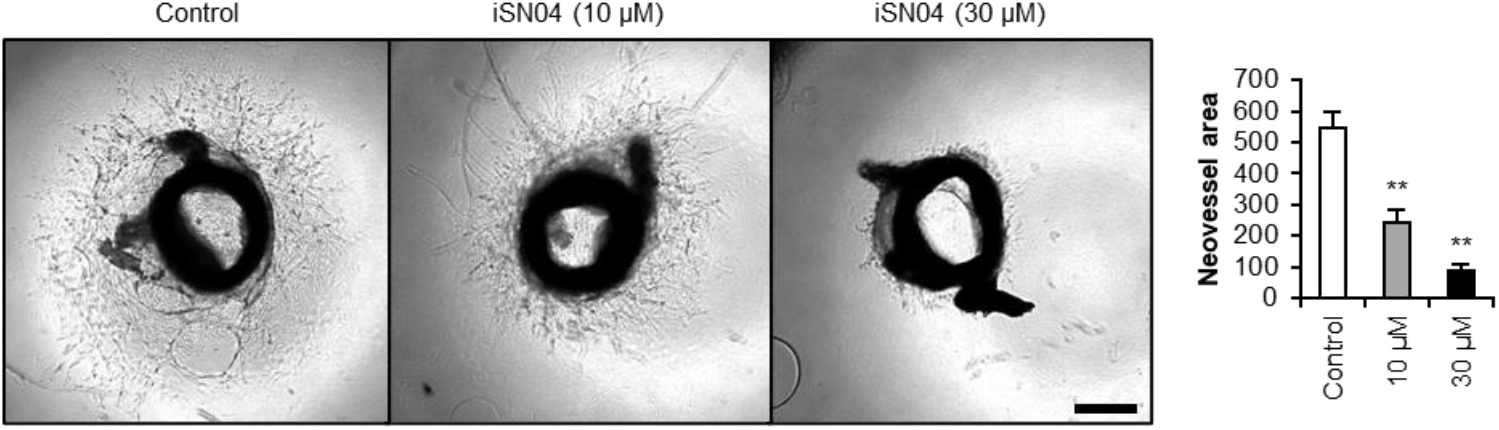
iSN04 suppresses angiogenesis of aortic rings. Representative microscopic images of aortic rings treated with 10 or 30 μM iSN04 in GM for 6 days. Scale bar, 200 μm. Neovascular sprouting area was quantified. ** *p* < 0.01 vs control (Scheffe’s *F* test). *n* = 5-6.

## 4. Discussion

The present study demonstrated that the 18-base myogenetic anti-nucleolin aptamer, iSN04, suppresses proliferation and promotes differentiation of VSMCs, resulting in the inhibition of aortic ring angiogenesis. Nucleolin has been reported to promote proliferation and inhibit differentiation of VSMCs, and nucleolin knockdown reverses the cellular phenotypes [15-17]. Our results of nucleolin antagonism by iSN04 are well consistent with these findings and provide a novel biomolecule for nucleolin inhibition in VSMCs.

The mechanism of action of iSN04 in VSMCs needs to be further investigated for clinical application. The effect of iSN04 on VSMCs closely resembled to that on myoblasts [5]. The previous studies reported that antagonism of nucleolin by iSN04 or another anti-nucleolin aptamer, AS1411, increased p53 protein levels in myoblasts and glioma cells, respectively [5,30], because nucleolin normally binds to the 5’ untranslated region of p53 mRNA to inhibit its translation [31,32]. p53 is known to be required for SMC differentiation [33]. SMC-specific overexpression of p53 suppresses injury-induced neointima formation [33] and endogenous levels of p53 protect VSMCs from apoptosis and prevent plaque formation [34]. These findings suggest that iSN04-induced p53 protein levels may be part of the induction of VSMC differentiation.

In addition, our previous study reported that iSN04 suppresses tumor necrosis factor-α (TNF-α)-induced inflammatory responses of skeletal and smooth muscle cells [12]. Since nucleolin interferes with the degradation of β-catenin through Ser9 phosphorylation of glycogen synthase kinase 3β (GSK3β) [35], nucleolin inhibition by iSN04 reduces nuclear accumulation of β-catenin with nuclear factor-κB (NF-κB), resulting in the suppression of inflammatory transcription [12]. β-catenin is also induced in VSMCs after injury to facilitate neointima formation, and β-catenin inhibitors prevents VSMC proliferation [36]. Thus, the decrease in β-catenin protein levels by iSN04 may be another mechanism for the inhibition of VSMC growth.

Switching de-differentiated VSMCs to a differentiated state is a key strategy for the prevention and treatment of widespread vascular diseases such as atherosclerosis [2], neointima formation after injury and during restenosis [4], and pathological angiogenesis in atherosclerosis, arthritis, diabetic retinopathy, or cancer [29]. iSN04 is the only 18-base anti-nucleolin aptamer that can be chemically synthesized and modified on a large scale at low cost, which would be suitable for a nucleic acid drug [37]. The treatment of vascular diseases with nucleic acid aptamers has been studied. Pegaptanib (Macugen) is the antivascular endothelial growth factor (VEGF) 165 aptamer for neovascular age-related macular degeneration and the first therapeutic aptamer approved by the U.S. Food and Drug Administration (FDA) in 2004. However, pegaptanib is now rarely used because later anti-VEGF antibodies such as ranibizumab or soluble VEGF receptors such as aflibercept are more effective [38]. As of the end of 2023, the FDA has approved only two aptamers; pegaptanib and avacincaptad pegol (Izervay), the anti-C5 complement aptamer for geographic atrophy. The development of novel candidates is an important issue to realize aptamer therapy in clinical settings. The present study demonstrated that the anti-nucleolin aptamer iSN04 may be useful for the treatment of vascular diseases. Its efficacy and safety should be further investigated in animal models in future studies.

## 5. Conclusions

An 18-base myogenic anti-nucleolin aptamer, iSN04, suppressed proliferation and promoted differentiation of VSMCs, resulting in inhibition of aortic angiogenesis. iSN04 may be a nucleic acid drug to treat de-differentiated VSMCs in vascular diseases such as atherosclerosis, neointima formation, and pathological angiogenesis.

## Author Contributions

Conceptualization, T.T.; investigation, M.M. and T.T.; resources, T.S.; writing—original draft preparation, T.T.; funding acquisition, M.M. and T.T. All authors have read and agreed to the published version of the manuscript.

## Institutional Review Board Statement

The experimental procedure for the preparation of mouse aortic rings was performed in accordance with the Regulations for Animal Experimentation of Shinshu University, and the animal protocol was approved by the Committee for Animal Experiments of Shinshu University (No. 022076).

## Informed Consent Statement

Not applicable.

## Data Availability Statement

The data presented in this study are available on request to the corresponding author.

